# “Paraxenoviridae”, a putative family of ubiquitous marine bacteriophages with double-stranded RNA genomes

**DOI:** 10.1101/2025.02.27.640542

**Authors:** Mitsuhiro Yoshida, Sofia Medvedeva, Akihito Fukudome, Yuri I. Wolf, Syun-ichi Urayama, Yosuke Nishimura, Yoshihiro Takaki, Eugene V. Koonin, Mart Krupovic, Takuro Nunoura

## Abstract

<u>F</u>ragmented and primer <u>L</u>igated <u>D</u>sRNA <u>S</u>equencing (FLDS) was used to reconstruct five complete, bisegmented RNA genomes of paraxenoviruses, a group of viruses that was previously identified in the ocean and that based on the analysis of partial genomes was proposed to represent a putative new phylum within the kingdom *Orthornavirae* of the realm *Riboviria*. Phylogenetic analysis of the RNA-directed RNA polymerases of paraxenoviruses demonstrated their affinity with the ribovirus order *Durnavirales* within the class *Duplopiviricetes* of the phylum *Pisuviricota*. The order *Durnavirales* includes families *Cystoviridae* that consists of well-characterized dsRNA bacteriophages and less thoroughly studied *Picobirnaviridae* that are also suspected to infect bacteria. Consistently, modeling and analysis of the structure of the predicted capsid protein (CP) of several paraxenoviruses revealed similarity to picobirnavirus CP although the paraxenovirus CP is much larger and contains unique structural elaborations. Taken together, these affinities suggest that paraxenoviruses represent a distinct family within *Durnavirales*, which we provisionally name “Paraxenoviridae”. Both genomic segments in *Picobirnaviridae* and “Paraxenoviridae” encompass multiple open reading frames, each preceded by a typical bacterial ribosome-binding site, strongly suggesting that these families consist of bacterial viruses. Search for homologs of paraxenovirus genes shows widespread distribution of this virus group in the global ocean, suggesting a potential important contribution to marine microbial ecosystems. Our findings further expand the diversity and ecological role of the bacterial RNA virome, reveal extensive structural variability of RNA viral capsid proteins, and demonstrate the common ancestry of several distinct families of bacterial viruses with dsRNA genomes.

## Introduction

In the comprehensive taxonomy of viruses that was recently adopted by the International Committee on Taxonomy of Viruses (ICTV), RNA viruses without a DNA stage in their reproduction cycles comprise the kingdom *Orthornavirae* within the realm *Riboviria* [1]. All these viruses share a single conserved gene that encodes the enzyme responsible for virus RNA replication, RNA-directed RNA polymerase (RdRP). Based on the phylogeny of the RdRPs, the kingdom *Orthornavirae* splits into 7 phyla corresponding to 5 major clades in the tree, *Lenarviricota*, *Pisuviricota*, *Kitrinoviricota*, *Duplornaviricota*, and *Negarnaviricota* [2–4], and 2 smaller phyla, *Ambiviricota* and *Artimaviricota*, created more recently [5, 6].

In the last few years, culture-independent metatranscriptome analysis followed by phylogenetic analyses of RdRPs dramatically expanded the diversity and ecological spread of RNA viruses in the kingdom *Orthornavirae* [7–11]. For example, metatranscriptome analysis of a single large coastal water sample from the Yangtze estuary in China yielded more than 4,500 previously unidentified distinct members of *Orthornavirae*, doubling the number of known RNA virus lineages at the level between species and genus [8]. Subsequent, large scale metatranscriptome studies have expanded the diversity of the known riboviruses by at least an order of magnitude [9–11]. Most of the discovered viruses fall into the original 5 phyla, but in the latest Global Ocean RNA metatranscriptome study, 5 new candidate phyla have been proposed (*Taraviricota*, *Pomiviricota*, *Arctiviricota*, *Paraxenoviricota*, and *Wamoviricota*) [11], and two distinct candidate phyla were proposed in another large scale metatrancriptome study [10]. More recently, an artificial intelligence approach was employed to discover about 162,000 potential RNA virus species and 180 RNA virus supergroups from more than 10,000 global metatranscriptomes [12].

The recent updates of the diversity of RNA viromes are generally based only on RdRP sequence whereas complete genomes are necessary to infer with confidence virus replication and expression strategies, and to elucidate evolutionary relationships. This a challenging task, particularly, in the case of viruses with multi-segmented genomes because genes encoding proteins other than the RdRP are not highly conserved among RNA viruses [13]. To overcome this problem, adequate methods are required to obtain complete RNA virus genome sequences. We developed a dsRNA-derived cDNA construction and sequencing method, FLDS (<u>F</u>ragmented and primer <u>L</u>igated <u>D</u>sRNA <u>S</u>equencing), that enabled us to determine complete sequences of intracellular long dsRNA molecules [14, 15], such as genomes of dsRNA viruses and replication intermediates of ssRNA viruses [16]. FLDS also enables reconstruction of complete genomes for multi-segmented RNA viruses, given the similarities among the respective terminal sequences [14, 15, 17, 18]. Recently, FLDS has been successfully applied to characterize complete segmented genomes of two novel groups of bacterial riboviruses, one of which represents the new phylum, *Artimaviricota*, that might be subsequently upgraded to the kingdom rank [6].

‘*Candidatus* (*Ca.*) Paraxenoviricota’ has been proposed as a new phylum within the kingdom *Orthornavirae* based on the phylogenetic placement of a limited number of RdRP sequences from a global ocean metatranscriptome [11]. In the RdRP tree of Zayed *et al.* [11], paraxenoviruses form a distinct clade outside the phylum *Kitrinoviricota*. However, Edgar [19] presented an alternative phylogenetic analysis in which paraxenoviruses are lodged within *Kitrinoviricota*, leaving uncertain the phylogenetic and taxonomic positions of this group of viruses. Furthermore, no complete genomes of paraxenoviruses have been reported.

We employed FLDS to reconstruct complete bi-segmented genomes of paraxenoviruses from RNA viromes of pelagic surface water microbial communities from the North Pacific and East Indian Oceans. The expanded dataset including previously uncharacterized paraxenovirus lineages, allowed us to reassess the phylogeny of their RdRPs which showed that paraxenoviruses are related to members of the order *Durnavirales* within the class *Durnaviricetes* of the phylum *Pisuviricota*. We also identified the capsid protein (CP) of paraxenoviruses that has an elaborate predicted structure distantly related to that of the CPs of picobirnaviruses, one of the families of *Durnavirales.* Taken together, these findings further increase the diversity of the bacterial RNA virome that was expanded through the recent effort in metatranscriptome mining, reveal previously unknown structural variability of capsid proteins of RNA viruses with dsRNA genome, and demonstrate common origin of several groups of these viruses. We further demonstrate that paraxenoviruses are widely represented across the global ocean, suggesting a potential impact of these dsRNA bacterial viruses on marine microbial ecosystems.

## Materials and methods

### Sample collection

Four pelagic surface water samples, which ‘*Ca.* Paraxenoviricota’ sequences were obtained, were collected from each two stations in the North Pacific and the East Indian Ocean during JAMSTEC cruises (Table 1). At each station, approximately 10 L of surface water was collected by a bucket, and each 3–4 L of seawater was filtered with a 0.2-μm-pore-size cellulose acetate filter (Advantec, Tokyo, Japan). The filters were stored at –80°C until nucleic acid extraction as described previously [15].

**Table 1.** Property of the pelagic surface water samples.

| Library name | Curise ID | Station ID | Geographical coordination | Area | Sampling date |
| --- | --- | --- | --- | --- | --- |
| UraH2 | KM17-01 | Stn 8 | 29°18' N,<br>143°31' E | Izu-Ogasawara Trench, Northwest Pacific Ocean | 06-Jan-2017 |
| UraH6 | MR15-05 | Stn 22 | 18°32' S,<br>111°52' E | East Indian Ocean | 01-Jan-2016 |
| UraH20 | MR14-04 | Stn 147 | 47°0' N, 126°<br>0' W | Northeast Pacific Ocean | 22-Aug-2014 |
| UraH22 | MR17-03C | 005 (SC4/KG5) | 25°30' N,<br>126°5' E | Okinawa Trough, East China Sea | 31-May-2017 |

### RNA extraction

The RNA extraction method was described previously [15]. Cells collected on a portion of the 0.2-μm-pore-size filters corresponding to approximately 2 L of seawater were pulverized in a mortar in liquid nitrogen and suspended in dsRNA extraction buffer [20 mM Tris–HCl, pH 6.8; 200 mM NaCl; 2 mM EDTA; 1% SDS; 0.1% (v/v) β-mercaptoethanol]. The manually extracted total dsRNA using an SDS-phenol method was further purified with the cellulose resin chromatography [20, 21] as described previously [15].

### cDNA synthesis and Illumina sequencing library construction and sequencing

Libraries for FLDS were synthesized from the extracted total RNA samples [15, 18]. In brief, the dsRNA was physically fragmented into about 1.5 kbp and ligated an adapter oligonucleotide to 3’-end of fragmented dsRNAs. After heat denaturation with an oligonucleotide primer, which consisted of complementary sequence to the adapter oligonucleotide, cDNA was synthesized using SMARTer RACE 5’/3’ Kit (Takara Bio, Shiga, Japan). After PCR amplification, short DNA fragments including the primers were removed using an 80% volume of AMPure XP (Beckman Coulter, Brea, CA, United States), and the purified cDNA was fragmented using the ultrasonicator Covaris S220 (Covaris, MA, USA).

Illumina sequencing libraries (Table S1) were constructed using KAPA Hyper Prep Kit Illumina platforms (Kapa Biosystems, MA, USA) from the physically sheared cDNAs constructed from the dsRNA. The libraries were sequenced using the Illumina MiSeq v3 Reagent Kit (600 cycles) with 300-bp paired-end reads on the Illumina MiSeq platform.

### Data processing for RNA virome community analysis and viral genome reconstruction

The raw reads from the dsRNA-derived cDNA libraries (Table S1) were processed with a custom Perl script (https://github.com/takakiy/FLDS) [17]. In brief, Illumina adaptor sequences, cDNA synthesis adaptors, and low-quality or low-complexity reads were trimmed from the obtained raw reads as previously described [15, 17]. Reads of rRNA sequences were removed using SortMeRNA 2.0 [22]. Cleaned-up reads were subjected to *de novo* assembly using CLC Genomics Workbench ver. 21.0 (Qiagen, Tokyo, Japan) with the following parameters: a minimum contig length of 500, word value set to 39, and bubble size set to 300.

To analyze the composition of the pelagic RNA virome communities, RdRP coding contigs were scanned from the assembles of each sample using HMM search with several RdRP HMM profiles, including Pfam 34.0 [23], RdRP hmm profiles used by Zayed *et al.* [11], and NeoRdRp program [24]. All predicted RdRP contigs were clustered at 96% similarity using CD-HIT-EST [25]. Cleaned-up reads of each library were mapped against representative contigs (i.e., the longest contigs) in the clusters. FPKM (<u>F</u>ragments <u>p</u>er <u>K</u>ilobase <u>M</u>illion mapped reads) of each mapped virus contig was calculated form the associated read counts.

In order to reconstruct complete genomes of multi-segmented RNA viruses from the contigs assembled in this study, a custom Perl script (https://github.com/takakiy/FLDS) was used to determine terminal sequences of RNA viral genome segments [17]. The completeness of terminal sequences of each RNA viral genome segment was determined based on the following cases: (i) reads with adaptor sequence were aligned at both termini of a genomic segment, and (ii) their frequencies are generally higher than those in the central region of the segment. The grouping of full-length genomic segments derived from a multipartite RNA virus or its population was conducted based on the conserved sequences at both termini of these segments [6, 14, 15, 17, 18]. The open reading frames (ORFs) and their upstream ribosome-binding sites (RBSs) were identified using Prodigal version 2.6.3 [26] and by manual inspection. The identification and taxonomic assignment of RdRP genes in these virus genome candidates were examined using the HMM searches with Pfam, RdRP hmm profiles of Zayed *et al.* [11], and NeoRdRp, as mentioned elsewhere. Transmembrane domains were predicted using TMHMM [27].

### Analysis of untranslated regions on the RNA virus genomes

To verify that the 5’ termini of RdRP-encoding segments of paraxenovirus genomes do not contain any conserved ORFs, the gene calling was performed with Prodigal using non-standard translational tables (1-25). The RNA secondary structures were predicted using ViennaRNA package with default parameters [28]. Trinucleotide frequency analysis was done in R (trinucleotideFrequency, library Biostrings) [29].

### Phylogenetic analysis

To identify additional RdRP sequences related to paraxenoviruses, a BLASTP search using paraxenovirus sequences obtained from this study and TARA study [11] was conducted against IMG/VR Viral Protein Database (IMG/VR v4: [30]) (*E*-value threshold: 1e-10). This search led to the identification of 41 RdRP sequences with high similarity to those of paraxenoviruses. Including these sequences, 48 core sequences of paraxenovirus full-length RdRPs were aligned using MUSCLE5 [31]; the alignment was used as a query in an HHSEARCH [32], which is run against a database of family-level RdRP core alignments from Neri *et al.* [10]. Because clusters of the order *Durnavirales* were identified as the closest relatives of paraxenoviruses, consensus sequences of 24 family-level clusters of *Durnavirales*, and one outgroup family (f.0121 of Neri *et al.* [10], that formed a sister branch to *Durnavirales* proper) were aligned together with the consensus of the 48 paraxenovirus RdRPs using MUSCLE5. Then, each consensus amino acid was expanded to the full alignment column, resulting in a complete alignment [10]. Alignment positions containing more than 67% gaps and with homogeneity below 0.05 were removed [33]. An approximate ML phylogenetic tree was reconstructed from the trimmed alignment of 4,453 sequences using the FastTree program with WAG evolutionary model and gamma-distributed site rates [34]. A set of up to 40 most diverse representatives was selected from each family-level alignment, forming a subset of 507 sequences with all families represented. An alignment of these representatives was extracted from the complete alignment, and a ML tree was reconstructed for this subset using IQ-TREE [35] with Q.pfam+F+R8 evolutionary model, selected by ModelFinder, and aBayes branch support values.

### Modeling of RdRP structures with AlphaFold2 and structural comparisons

To model paraxenovirus RdRP structures, for each prediction, we used Clustal Omega [36] to create a custom multiple sequence alignment (MSA), consisting of 5 sequences from this study (GT1–5), 11 sequences from the TARA study, and additional sequences from the IMG/VR database. Each of GT4, GT3, and GT5 RdRP sequences was used as a query for BLASTP search against IMG/VR Viral Protein Database [30] (*E*-value threshold: 1e-10), and highly similar homologs were identified (18, 24, and 13 sequences, respectively). TARA_132 RdRP model prediction used a custom MSA with the same sequence set as GT4 because BLASTP search against IMG/VR using TARA_132 RdRP sequence resulted in the same hits. All predictions were performed using locally installed colabfold 1.5.1 (--model-type alphafold2_ptm,--num-recycle 10), and the final models were AMBER-relaxed [37].

### Structural modeling of capsid proteins

To model the paraxenovirus capsid protein (CP) structure, we retrieved additional CP sequences from the LucaProt dataset [12] with an iterative profile search. One iteration included hmmsearch (-E e˄-10), the alignment of hits with MAFFT (--auto) [38] and construction of a new HMM profile for the next iteration using HMMER 3.4 [39]. The search was initiated using a profile created from the seed MSA of CPs from this study (GT1–5) and 11 CP sequences from IMG/VR database. After eight iterations, no more new sequences were found. Partial CPs sequences (less than 900 aa in length) were removed (seqtk seq-L 900).

Redundant sequences were filtered out with mmseqs2 (id 1, coverage 0.8). The final CP alignment used for modelling consists of 145 non-redundant sequences. In the modelling job, the CP sequence of GT2 was used as a representative query for GT1, GT2, and GT3. The prediction of dimers was performed using a local installation of ColabFold 1.5.3 (--model-type alphafold2_multimer_v3,--num-recycle 12).

## Results

### dsRNA Sequencing and identification of RdRPs of‘*Ca.* Paraxenoviricota’

The FLDS method enables identification of full-length viral genomic segments based on the read mapping of contigs, and complete sets of the genomic segments of multipartite RNA viruses can be identified based on the conserved sequences at the termini of contigs [14, 17]. Using this method, we sequenced 18 dsRNA metatranscriptomic libraries from North Pacific and Indian Oceans [15] (Yoshida *et al.*, in preparation). The datasets from each library were then searched for the presence of RdRPs with public and environmental RdRP Hidden Markov Model (HMM) profiles [11]. These searches yielded four full-length segments (SEG1 to SEG4) encoding ‘*Ca.* Paraxenoviricota’-type RdRPs from the FLDS-derived UraH2, UraH20, and UraH22 cDNA libraries (Table 1 and Table S2). A subsequent BLASTX search using the RdRP sequences of the newly identified SEG1–4 as the queries allowed the identification of the SEG5 segment from the UraH6 cDNA library (Table 1 and Table S2).

### Complete genomes of‘*Ca.* Paraxenoviricota’

Given that segments of the same virus genome typically share terminal sequences (e.g., [14, 15, 40]), we searched the FLDS libraries for the presence of segments with terminal sequences similar to those of the RdRP-encoding SEG1-5 segments. For each of the RdRP-encoding segments, we identified an additional putative genome segment harboring one or more open reading frames (ORFs) (Table S2). Accordingly, we concluded that the ‘*Ca*.

Paraxenoviricota’ viruses present in our datasets have bipartite genomes, which we refer to as GT1–5 (Genome ID). For each genome set, we designated the segment that harbored the RdRP-encoding ORF as RNA1 and the second segment that harbored non-RdRP ORFs as RNA2 (Fig. 1).

**Fig. 1.**
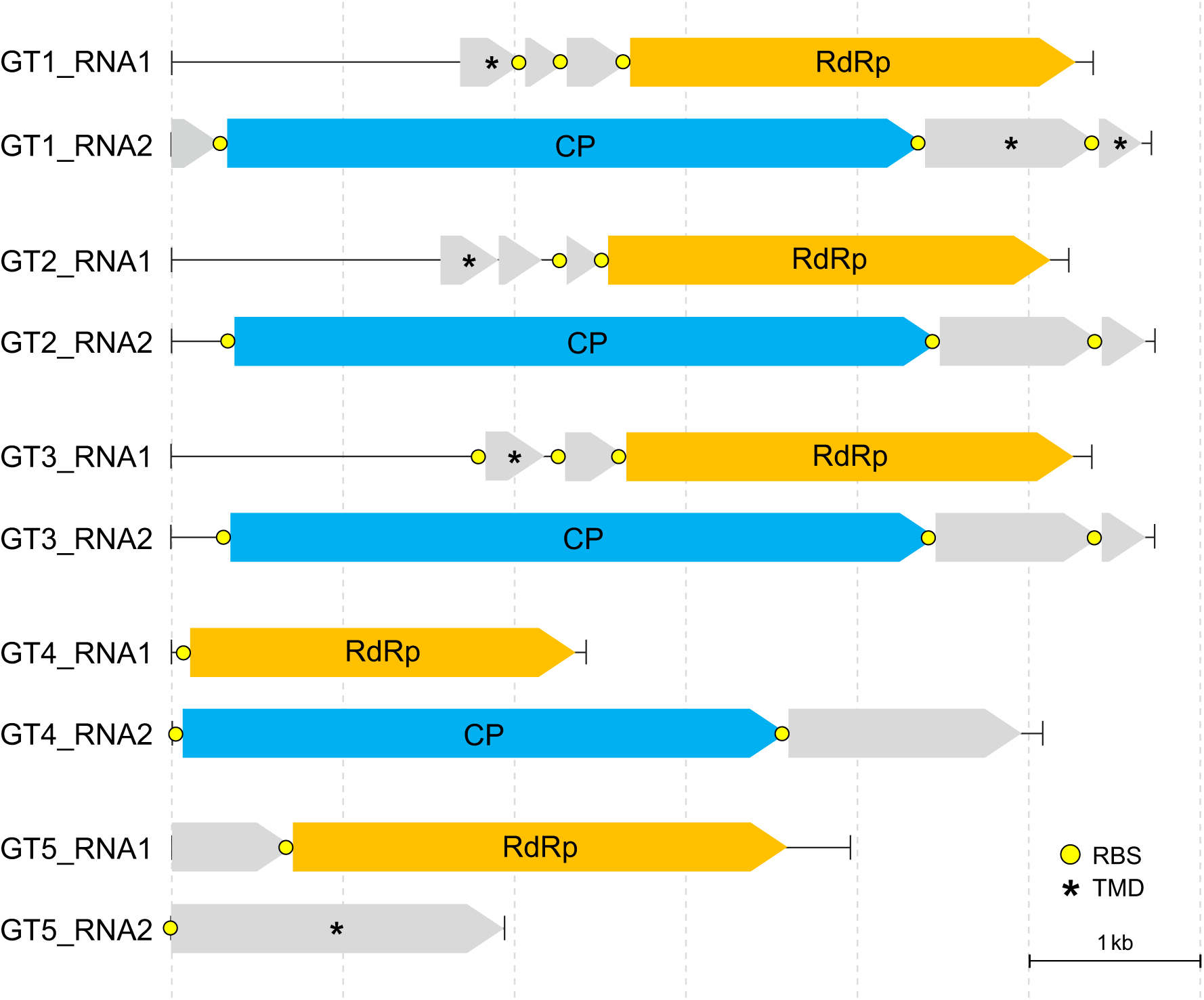
**Novel bipartite ‘*Ca.* Paraxenoviricota’-associated RNA virus genomes reconstructed from pelagic dsRNA virome samples**. Each of the identified five ‘*Ca.* Paraxenoviricota’-associated virus genomes consist of RNA1 and RNA2 genomic segments. Open reading frames encoding homologous RdRP proteins are shown as arrows with orange color. Open reading frames encoding homologous CP proteins are shown as arrows with light blue color. Yellow circles represent predicted SD RBS motifs. Asterisks denote putative genes encoding predicted transmembrane domain (TMD)-containing proteins.

RNA1 and RNA2 each harbored 1 to 4 ORFs encoding predicted proteins that varied in length from 80 to 1,376 amino acid residues (Fig. 1 and Table S3). All RNA1 segments shared the RdRP-encoding ORF, and in addition, GT1–3 encompassed two homolgous ORFs (Fig. S1). In RNA2 segments, three ORFs were found to be homologous between among GT1–3 (Fig. S2). The predicted paraxenovirus proteins showed no significant sequence similarity (*E*-value < 10^-4^) to any proteins in the non-redundant protein sequence database (NCBI) that was searched using BLASTP, or in the PFAM, CDD, and PDB databases that were searched using HHpred with the multiple alignments of the respective predicted paraxenovirus proteins used as queries (Probability > 50). However, we found that RNA1 of GT1–3 and RNA2 of GT1 and-5 harbored at least one ORF encoding a protein with a predicted transmembrane domain (Fig. 1), which could be involved in virus-host interactions [41]. The RdRP sequences of paraxenoviruses identified in this study showed the highest similarity to those encoded by the two contigs among 7 paraxenovirus contigs (Fig. S3 and Table S3) derived from the TARA RNA virome dataset of the Pacific Ocean [11]. Notably, most of the ORFs are preceded by typical bacterial-type RBSs (Shine-Dalgarno motifs [SD motifs]) (Fig. 1 and Table S3).

The genomic sequences of paraxenoviruses contain extended 5’-terminal regions that lack any long and/or conserved ORFs that could be predicted using the standard or any of the available alternative translation tables. These regions have markedly different trinucleotide compositions compared to the predicted protein-coding genes, in particular, those encoding the predicted RdRPs (Fig. 2C), strongly suggesting that the long 5’UTRs are indeed non-coding. Long UTRs are not uncommon in RNA viruses, but the ∼1.5 kb 5’UTRs of paraxenoviruses appear to be among the longest. Notably, although most likely non-coding, the nucleotide sequences of the 5’UTRs are conserved, especially between GT2 and GT3.

**Fig. 2.**
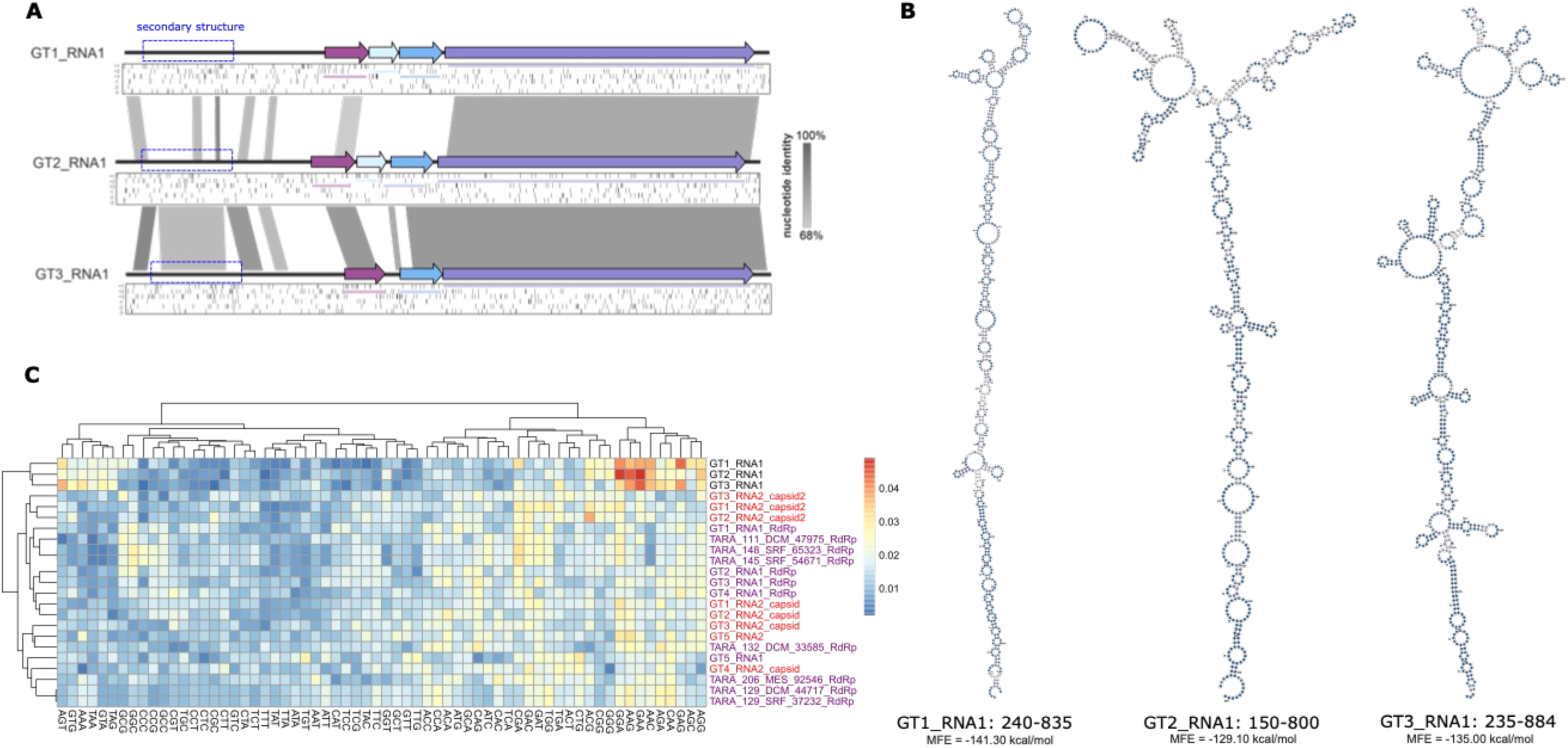
Long 5’ untranslated regions in paraxenovirus RNA segments. (A) Distribution of stop codons in GT1_RNA1, GT2_RNA1, and T3_RNA1. The positions of stop codons in all six translation frames are shown as black stripes below the genome map. Regions conserved between ree viruses (nucleotide identity > 68%) are connected by grey links. The dashed box shows a position of predicted secondary RNA structure. (B) redicted RNA secondary structure found in conserved non-coding regions of paraxenoviruses. The coordinates of secondary structures are indicated. C) Trinucleotide composition of paraxenovirus genomes. Frequency of trinucleotides in non-coding regions (top three rows of the heatmap) are ompared with CP (names in red) and RdRP (names in purple) coding regions.

Folding of these sequences suggests that the conserved regions (blue boxes in Fig. 2A) adopt similar secondary structures in all three genomic segments (see Fig. 2B). The UTRs might be involved in regulation of translation as, for example, shown for the 5’UTRs of picornaviruses [42, 43].

### Phylogenetic analysis of paraxenovirus RdRPs

Altogether, we identified 48 unique sequences of paraxenovirus RdRPs, which were trimmed to 40 maximally diverse representatives using a large set of full-length RdRp core sequences for the purpose of phylogenetic analysis [10]. In the maximum likelihood phylogenetic tree of the core sequences of orthornavirus RdRPs, paraxenoviruses were nested within the *Durnavirales* order of the *Duplopiviricetes* class in the phylum *Pisuviricota* (Fig.3). The sister group of paraxenoviruses is a yet unnamed group that was previously designated f.0117_base-cysto [10]. The clade comprising these two groups is lodged deeply within the *Durnavirales* branch of *Duplopiviricetes* and is supported by high bootstrap values (Fig. 3). Thus, there is a discrepancy between our findings and the previous placement of paraxenoviruses in the phylogeny of the *Riboviria* where these viruses were assigned to a putative new phylum [11]. The difference in the positions of paraxenoviruses in the two phylogenies might be due to the substantially smaller number of sequences from this group and related groups in the analysis of Zayed *et al.* [11].

**Fig. 3.**
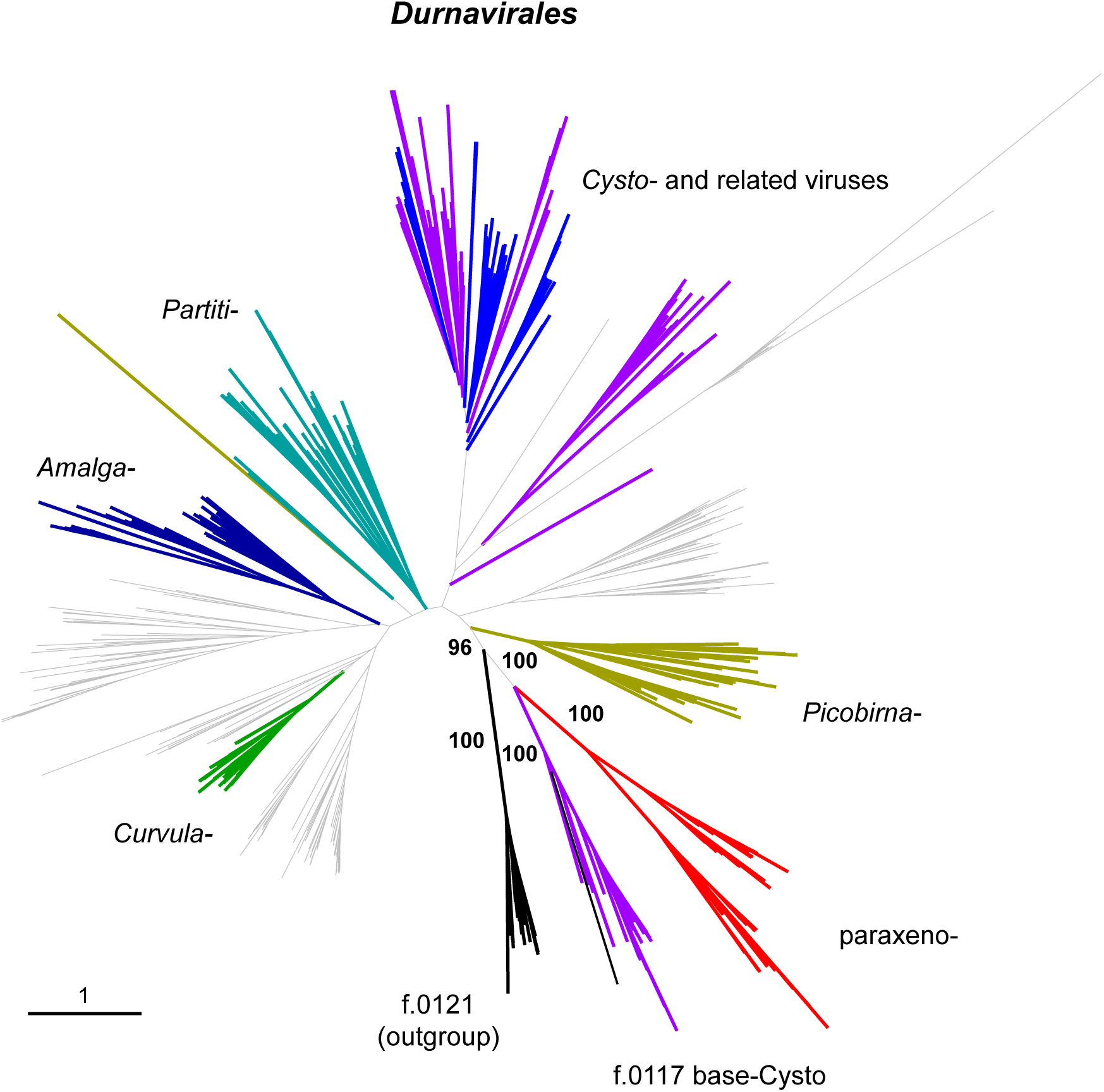
Phylogenetic tree of RdRPs sequences from paraxenoviruses and related viruses. A Maximum Likelihood tree (IQ-TREE, Q.pfam+F+R8 evolutionary model) was reconstructed using an alignment of RdRp core sequences of 507 representatives of *Durnavirales*, paraxenoviruses, and an outgroup family. Branches highlighted in black: f.0121 (outgroup); red: paraxenoviruses; light blue: *Cystoviridae*; magenta: basal cystovirus-like families; yellow: *Picobirnaviridae*; cyan: *Partitiviridae*; dark blue: *Amalgaviridae*; green: *Curvulaviridae*. aBayes support values are shown for the branches around paraxenoviruses.

Within the paraxenovirus group itself, the phylogeny splits into four clades (Fig. S4). The RdRPs of GT1,-2, and-3 formed a distinct subgroup in clade III, the GT5 RdRP belonged to clade IV, and the GT4 RdRP clustered with several TARA RdRPs (Clade I). For the following RdRP structure modeling, the RdRP sequences of GT3,-4, and-5 were used as representatives of clades III, I, and IV, respectively, and for clade II, TARA132 RdRP was selected.

### Structural modeling of paraxenovirus RdRPs

To further characterize paraxenoviruses, we predicted the structures of the newly identified RdRPs in GT3, GT4, and GT5 genomes along with one identified in the previous study (TARA132). The modeled structures of the paraxenovirus RdRPs were used as queries to search the BFVD database [44] of predicted viral protein structures and PDB database using FoldSeek. The RdRP structures of picobirnaviruses and partitiviruses were consistently identified as the most similar, with the structure of Mongoose picobirna-like virus [45] being the best hit with the probability of 1 (Tables S4 and S5). This result corroborates the phylogenetic placement of paraxenoviruses within *Durnavirales*. Thus, given the results of phylogenetic analysis (Fig. 3) and the structure-based searches, the predicted structural models of the paraxenoviral RdRPs were compared to those of structurally characterized durnaviral RdRPs, namely RdRPs of cystoviruses phi6 and human picobirnavirus, confirming the close similarity between paraxenovirus and picobirnavirus RdRPs (Fig. 4).

**Fig. 4.**
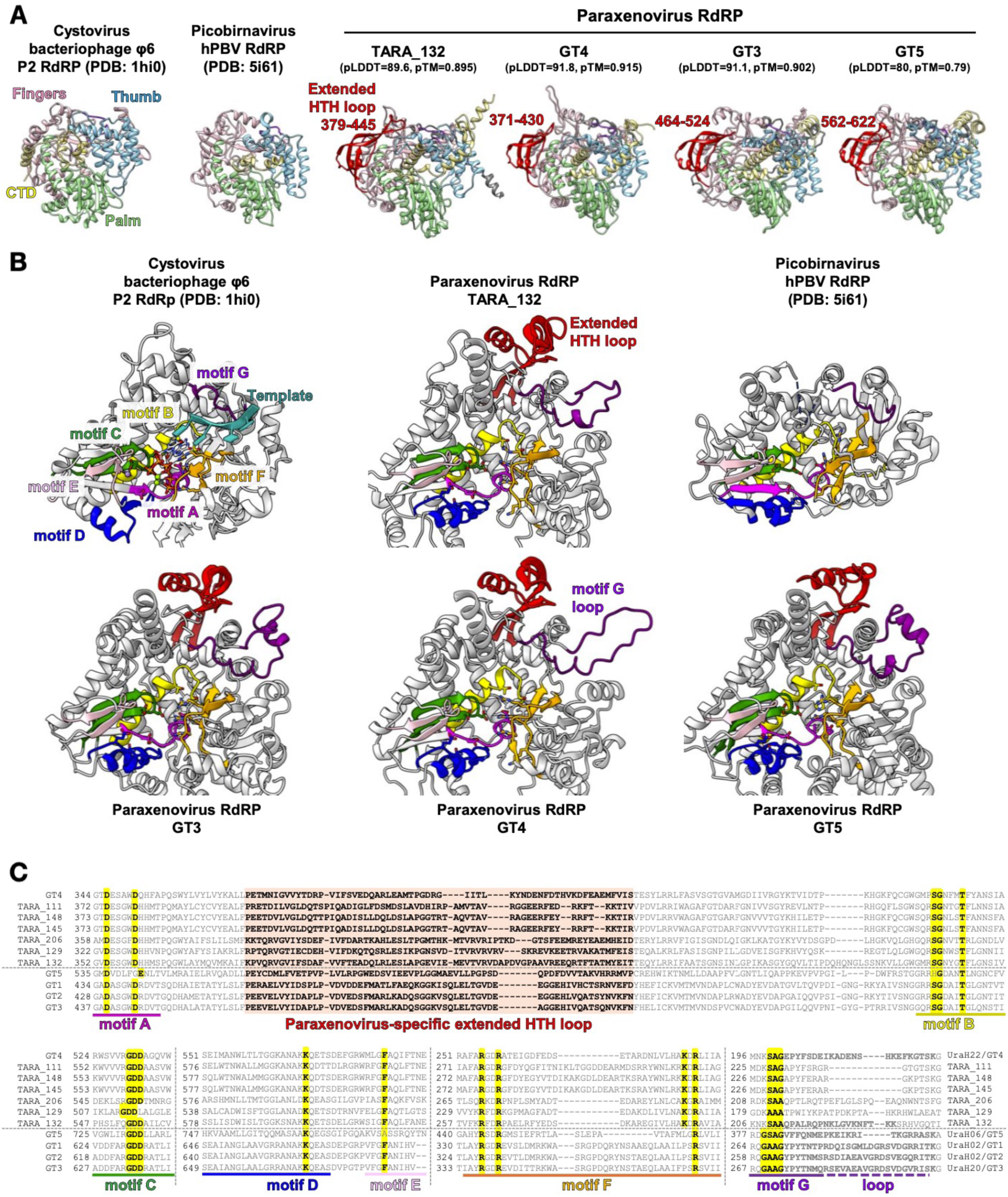
Predicted models and structural features of paraxenovirus RdRPs. (A) Subdomain comparison of AlphaFold-predicted paraxenovirus RdRP models with experimentally determined cystovirus and picobirnavirus RdRP structures. Residues 1–158 are hidden in the GT5 model due to low confidence. For the AlphaFold models, pLDDT (predicted local distance difference test) and pTM (predicted template modelling) quality scores are indicated. (B) Comparison of motifs A–G in the predicted paraxenovirus RdRP models to those in cystovirus and picobirnavirus RdRPs. Conserved signature residues in each motif are displayed in stick style. Thumb subdomain residues are hidden for clarity. For reference, Mg^2+^ (green dots), Mn^2+^ (a purple dot), two initiating GTPs and a template strand in the phi6 RdRP initiation complex are also shown (PDB:1hi0; [58]). (C) Amino acid sequence alignments of paraxenovirus RdRPs for inferred motifs A–G regions. Conserved signature residues in each motif and the extended HTH loop region are highlighted in yellow and orange, respectively.

As expected, the paraxenovirus RdRP models showed the canonical “right-hand” RdRP architecture consisting of the fingers-palm-thumb subdomains. All the models of paraxenovirus RdRPs contained a unique, extended, about 60 aa long insertion in a conserved helix-turn-helix (HTH) motif immediately after motif A (Fig. 4A). The insertion extended directly from the first α-helix of the HTH motif and contained multiple β-strands and an α-helix, joining back to the second α-helix of the HTH motif followed by a conserved β-hairpin (“middle” finger). Although insertion of helices and loops within the β-hairpin motif was detected in *Negarnaviricota* RdRPs, the insertion of a long loop within the HTH motif, extending toward a dsRNA exit channel, appears to be unique to paraxenoviruses and may serve as a defining feature of this virus group.

In addition to the core motifs A–C (Fig. 4A), other canonical RdRP motifs D–G can be inferred based on conservation of key residues in each motif and their spatial positions in the predicted models of paraxenovirus RdRPs, in comparison to publicly available structures of RdRPs from relatively close groups within *Durnavirales* such as cystoviruses and picobirnaviruses (Fig. 4B and 4C). Paraxenovirus RdRP motif G appears unique in containing a longer loop, following a canonical template-interacting region that encompasses conserved small residues such as serine, alanine and glycine [46]. The additional loop size varies among paraxenoviruses, as can be seen in sequences of TARA-associated RdRPs (Fig. 4C). Motif F of paraxenovirus RdRP is similar to that of cystovirus and picobirnavirus RdRPs, containing multiple positively charged arginine and lysine residues conserved in positions where they would interact with incoming NTPs. Based on motif F, paraxenovirus RdRPs can be divided into two subgroups, the first one containing three conserved arginines (GT1–3 and GT5) and the second one containing an additional, conserved lysine residue (GT4 and TARA-associated). In the former group, GT5 RdRP stands out further due to (1) the presence of a glutamate as the second negatively charged residue at 1-aa shifted position in motif A, (2) a histidine replacing the conserved serine in motif B, and (3) lack of the conserved aromatic amino acid in motif E. The GT5 RdRP also contained an extended N-terminal region that could not be modelled confidently, likely due to insufficient coverage (aa 1–158, predicted local distance difference test [pLDDT] < 50).

### Identification and structural modeling of paraxenovirus capsid proteins

In an attempt to gain further insight into the functions of paraxenovirus proteins, we modeled and analyzed structures for all conserved ORF products. This analysis showed that homologous proteins encoded by the long ORF1 of RNA2 segments of paraxenoviruses GT1-3 represent the putative capsid protein (CP) (Fig. 1 and S2). For the corresponding protein of GT2 (1376 aa), a good quality (pLDDT=71.8) model was obtained. The protein consists of an unstructured N-terminal domain (NTD; residues 1–46) and two structured domains, the shell (S; residues 47–319, 369–679, and 1219–1376) and projection (P; residues 320–368 and 680– 1218) domains (Fig. 5A). The S domain is predicted to play the key role in the icosahedral capsid (shell) formation, the P domain points away from the capsid and is likely involved in interaction with the host, whereas the N-terminal region contains 7 positively charged arginine/lysine residues and is likely to bind to the genomic RNA in the capsid interior. The S domain was modeled with high confidence (pLDDT=89.5), whereas the pLDDT scores for the NTD and P domain were considerably lower (see Fig. S5A). The top hit in DALI search queried with the S domain alone was the picobirnavirus CP (PDB: 2VF1; [47]). The Z-score was relatively low (Z=2.4), but manual inspection of the two structural models showed that the two CPs had the same fold, strongly suggesting they are homologous. In particular, the two CPs share a core of three two-stranded β-sheets (Fig. 5B), surrounded by a network of α-helices, all within the S domain. Notably, the predicted CP of paraxenoviruses is far more complex than that of picobirnaviruses, with an especially elaborate P domain consisting of a twisted β-barrel extending ∼7 nm above the capsid surface (∼3 times higher than in picobirnavirus).

**Fig. 5.**
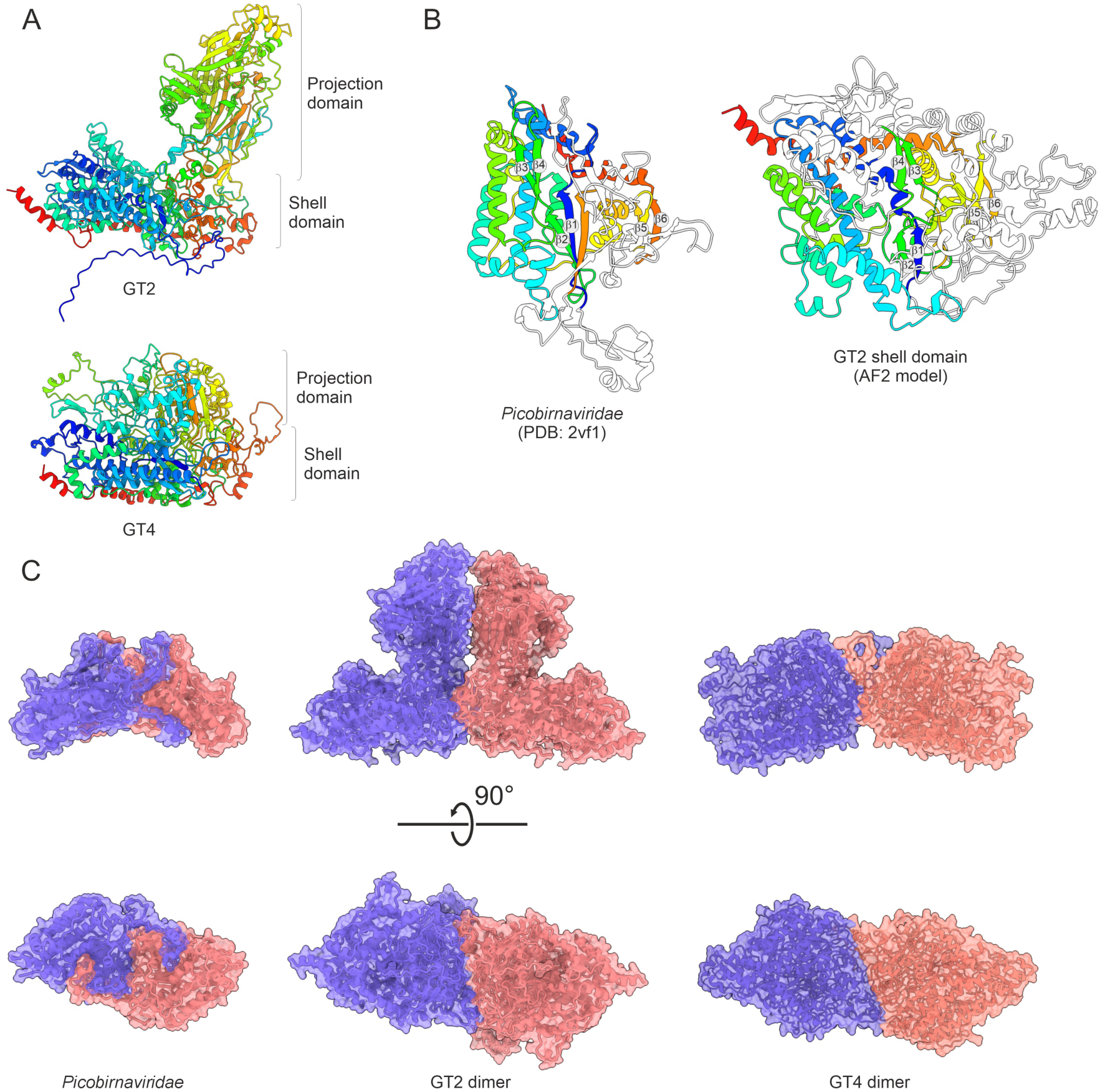
Predicted structures of capsid proteins of paraxenoviruses. (A) Domain organization of the paraxenovirus capsid proteins (CPs) encoded by GT2 (top) and GT4 (bottom). The structural models are colored using the rainbow scheme from blue N-terminus to red C-terminus. The shell and projection domains are labeled, whereas the unstructured N-terminal domain is omitted (refer to Fig. S5 for full model). (B) Comparison of the shell domains of the rabit picobinavirus (left) and paraxenovirus (right) CPs. In both models, the conserved structural elements are colored using the rainbow scheme, from blue N-terminus to red C-terminus, whereas the non-conserved elements are colored white. The six β-strands forming the three two stranded β-sheets are numbered. (C) Comparison of the CP dimers of rabbit picobinavirus (left, PDB id: 2vf1), paraxenoviruses GT2 (middle, pLDDT=70.8, ipTM [interface predicted template modelling]=0.661), and GT4 (right, pLDDT=75.2, ipTM=0.388). In all structures the subunits are colored red and blue. Side and bottom views are shown.

The protein (1177 aa) encoded by RNA2 of GT4 does not share recognizable sequence similarity with the CPs encoded by GT1–3. However, comparison of the corresponding structural models showed that the two proteins have the same fold (Fig. 5A). Notably, the similarity was restricted to the S domains, whereas the P domains were unrelated and the NTD was missing in the CP altogether. The P domain of GT4 CP did not form an extended structure elevated above the S domain. Nevertheless, similar to the S domain of GT2 CP, DALI search queried with the GT4 CP model as the query resulted in the best hit to the picobirnavirus CP (PDB: 2VF1) with a significant Z-score of 5.7.

Capsids of picobirnaviruses, similar to many other dsRNA viruses, including partitiviruses, cystoviruses, and totiviruses, are assembled from CP dimers, with 60 CP dimers forming a T=1 icosahedral capsid [47–51]. Therefore, we modeled the putative dimers of the paraxenovirus CPs. Although the predicted aligned error (pAE) for the P domain was relatively high, the confidently predicted dimerization interface within the S domain of GT2 involved regions equivalent to those involved in the picobirnavirus CP dimerization (Fig. 5C and S5D). A dimer prediction for GT4 also showed a similar dimerization pattern, though with lower confidence (Fig. 5C, ipTM [interface predicted template modelling]=0.388). This observation further supports the homology between the CPs of paraxenoviruses and picobirnaviruses.

### Relative abundance and biogeography of paraxenoviruses in marine habitats

We determined the community composition of the four pelagic intracellular RNA viromes from the North Pacific (UraH2,-20, and-22) and East Indian Ocean (UraH6), including the paraxenovirus populations, based on read abundance (FPKM) of the detected RdRP OTUs (Fig. 6A). In addition to paraxenoviruses, members of several ribovirus families with dsRNA genomes previously detected in marine habitats, including *Picobirnaviridae*, *Totiviridae*, and *Sedoreoviridae* [8, 15], were identified in these viromes. The relative abundance of paraxenoviruses varied from 2% to 18% of the RNA viromes.

**Fig. 6.**
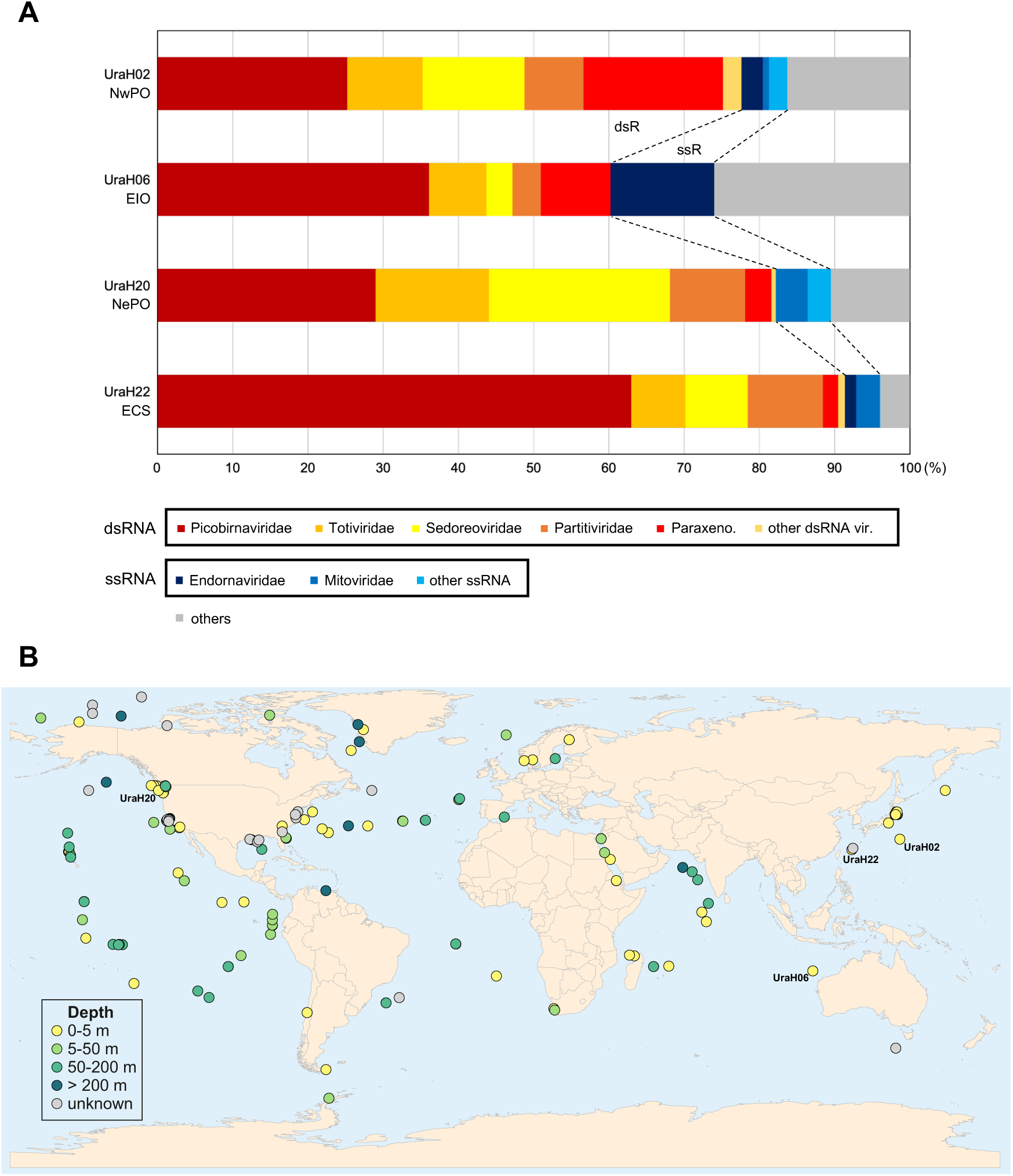
Distribution of paraxenoviruses in samples from the world ocean. (A) RNA virome community structures based on read abundance of RNA-dependent RNA polymerase (RdRp) phylotypes detected in this study. A family-level breakdown of the results is shown. The “other dsRNA” and “other ssRNA” categories represent their associated virus taxa that share less than 2% of the total sequences. The “others” category represents sequences of unclassified viruses. The biogeographic areas of the virome samples are represented as follows: NwPO (Northwest Pacific Ocean), EIO (East Indian Ocean), NePO (Northeast Pacific Ocean), and ECS (East China Sea). (B) Global distribution of paraxenoviral RdRP-and CP-encoding contigs in marine ecosystems. The circles represent the locations from which samples containing paraxenovirus-like sequences were collected. Circle colors indicate the water depth.

To analyze the ecological and geographic distribution of the paraxenovirus members, we assembled a dataset of orthologous RdRP (n=658) and CP (n=800) sequences from IMG/VR, LucaProt, and Tara Oceans databases for which information on the sampling sites was available. Analysis of the extracted metadata showed that paraxenoviruses are exclusive to marine ecosystems, are globally distributed, and are found in epipelagic and mesopelagic layers of the water column.

## Discussion

In this work, we sequenced five complete bipartite genomes of paraxenoviruses from pelagic surface waters. In contrast to the previous study that failed to reveal affinity between paraxenoviruses and any other ribovirus group [11], our analysis of the paraxenovirus proteins yielded strong indications that this virus group belongs to the class *Duplopiviricetes* and, probably, to the order *Durnavirales* within phylum *Pisuviricota*. The evidence of this affinity is threefold. First, in the RdRP phylogeny, paraxenoviruses, together with an uncharacterized related virus family, form a strongly supported clade within *Durnavirales* (Fig. 3). Second, comparison of the modeled structures of the paraxenovirus RdRPs confirmed the highest structural similarity with the RdRPs of picobirnaviruses, one of the families of *Durnavirales* (Table S4 and S5). We suspect that the previously reported lack of affinity of paraxenoviruses with any other riboviruses or alternative placement of this group within *Kitrinoviricota* was due to the relatively low representation of this group in the initially analyzed samples. Third, the structural model of the predicted paraxenovirus CP is similar to the solved structure of picobirnavirus CP although the paraxenovirus CP is much larger and has a far more elaborate structure than the picobirnavirus counterpart. Thus, paraxenoviruses appear to be relatives of picobirnaviruses, partitiviruses, and cystoviruses, and their most plausible taxonomic affiliation is a family within the order *Durnavirales*, for which we provisionally suggest the name “Paraxenoviridae”.

The apparent affinity of paraxenoviruses with *Durnavirales* has notable implications.

First, similarly to other viruses of this order, paraxenoviruses most likely have dsRNA genomes. Indeed, our analyses of pelagic RNA virome communities in this study and the previous study [15] revealed the presence of diverse dsRNA ribovirus families, further reinforcing the case for paraxenoviruses. The ecological underpinning of the diversity of the dsRNA virus genomes in these marine environments remains to be explored. Second, in each of the genomic segments of both picobirnaviruses and paraxenoviruses, the presence of multiple ORFs, each preceded by a typical RBS, strongly suggests bacterial hosts. Thus, the order *Durnavirales* seems to be dominated by bacteriophages with dsRNA genomes, although eukaryotic viruses infecting plants, fungi, and protists represent a sizable fraction as well [52]. More generally, these findings extend the trend that became prominent in recent metatranscriptome mining studies, namely, expansion of the diversity of RNA bacteriophages far beyond its previously perceived scope (e.g., [10]). Furthermore, our results reveal unprecedented variability of the capsid protein structures of viruses with dsRNA genomes.

These findings emphasize the potential of increased sequence coverage combined with detailed comparative genomic analysis to clarify the phylogenetic positions of virus families that initially appear unrelated to any known groups of viruses.

Until now, the information on dsRNA bacteriophages remained scarce due to a low number of isolates [53], and dsRNA phages of marine origin have not been identified. The results of the present work suggest the addition of the potential dsRNA “Paraxenoviridae” family to the growing diversity of known dsRNA phage groups, and furthermore, we showed that this family is widely distributed in different oceans, from polar to tropical regions. Given the presence of the cosmopolitan paraxenoviruses in marine ecosystems, further research may shed light on the host-virus relationships of these dsRNA phages (e.g., lytic versus non-lytic life cycle, chronic infection etc.) [54, 55], and consequent contributions to the viral shunt in the case of the lytic life cycle [56, 57]. Recently, marine metatranscriptome studies [10, 11] have also reported contigs of potential dsRNA phages within the phylum *Pisuviricota*. Thus, additional dsRNA phages likely exist in marine environments, and our FLDS-based approach can be expected to help unveiling environmental dsRNA viromes. Further FLDS-based RNA sequencing coupled with vertical and horizontal high-resolution sampling of pelagic waters is expected to provide insights into the diversity, ecology, and evolution of unexplored RNA viruses in the global ocean.

## Acknowledgements

We would like to thank the captain, crew, and onboard scientists and technicians of the R/V *Mirai* and R/V *Kaimei* of JAMSTEC during the MR14-04, MR15-05, and MR17-03C and KM17-01 cruises, respectively. We are grateful to Miho Hirai, Fumie Kondo, and Miwako Tsuda for technical assistance for RNA sequencing.

## Author contributions

M.Y. designed this work, analyzed the data, and wrote the manuscript. S.U. performed the experiments and wrote the manuscript. Y.T., Y.I.W., A.F., and S.M. analyzed the data and wrote the manuscript. Y.N. analyzed the data. T.N., E.V.K., and M.K. designed the work and wrote the manuscript.

## Conflicts of interest

JAMSTEC holds a patent for the ‘Double-stranded RNA fragmentation method and use thereof’, with S.U. and T.N. listed as inventors. These patents include European Patent (EP) Registration No. 3363898, registered on 30 November 2022; China Registration No.

ZL201680060127.X, registered on 8 February 2022; US Registration No. 10894981, registered on 19 January 2021; and Japanese patent No. 6386678, registered on 17 August 2018. The other authors declare no conflicts of interest.

## Funding

This work was supported in part by a Grant-in-Aid for Scientific Research (Grant No. 21K06312 to M.Y.) from the Japan Society for the Promotion of Science (JSPS), a Grant-in-Aid for Challenging Research (Grant No. 20K20377 to T.N.) from JSPS, and Grants-in-Aid for Scientific Research on Innovative Areas from the Ministry of Education, Culture, Science, Sports and Technology (MEXT) of Japan (Grant Nos. 17H05830 to M.Y.; 22H04879 and 20H05579 to S.U.; 19H05684, 16H06429, 16K21723, and 16H06437 to T.N.), and by Lilly Endowment, Inc., through its support for the Indiana University Pervasive Technology Institute which provided supercomputing resources for protein structure modelling. S.M. and M.K. were supported by Agence Nationale de la Recherche (grant ANR-23-CE02-0022).

Y.I.W. and E.V.K. were supported through the Intramural Research Program of the National Institutes of Health of the USA (National Library of Medicine).

## Supplementary data

Datasets of the segment sequences of five paraxenovirus genomes identified in this study are available at bioRxiv online.

